# The role of genetically distinct central amygdala neurons in appetitive and aversive responding assayed with a novel dual valence operant conditioning paradigm

**DOI:** 10.1101/2023.07.07.547979

**Authors:** Mariia Dorofeikova, Claire E. Stelly, Anh Duong, Samhita Basavanhalli, Erin Bean, Katherine Weissmuller, Natalia Sifnugel, Alexis Resendez, David M. Corey, Jeffrey G. Tasker, Jonathan P. Fadok

## Abstract

To survive, animals must meet their biological needs while simultaneously avoiding danger. However, the neurobiological basis of appetitive and aversive survival behaviors has historically been studied using separate behavioral tasks. While recent studies in mice have quantified appetitive and aversive conditioned responses simultaneously (Heinz et al., 2017; Jikomes et al., 2016), these tasks required different behavioral responses to each stimulus. As many brain regions involved in survival behavior process stimuli of opposite valence, we developed a paradigm in which mice perform the same response (nosepoke) to distinct auditory cues to obtain a rewarding outcome (palatable food) or avoid an aversive outcome (mild footshoock). This design allows for both within- and between-subject comparisons as animals respond to appetitive and aversive cues. The central nucleus of the amygdala (CeA) is implicated in the regulation of responses to stimuli of either valence. Considering its role in threat processing (Haubensak et al., 2010; Wilensky et al., 2006) and regulation of incentive salience (Warlow and Berridge, 2021), it is important to examine the contribution of the CeA to mechanisms potentially underlying comorbid dysregulation of avoidance and reward (Bolton et al., 2009; Sinha, 2008). Using this paradigm, we tested the role of two molecularly defined CeA subtypes previously linked to consummatory and defensive behaviors. Significant strain differences in the acquisition and performance of the task were observed. Bidirectional chemogenetic manipulation of CeA somatostatin (SOM) neurons altered motivation for reward and perseveration of reward-seeking responses on avoidance trials. Manipulation of corticotropin-releasing factor neurons (CRF) had no significant effect on food reward consumption, motivation, or task performance. This paradigm will facilitate investigations into the neuronal mechanisms controlling motivated behavior across valences.

**Significance Statement:** It is unclear how different neuronal populations contribute to reward- and aversion-driven behaviors within a subject. To address this question, we developed a novel behavioral paradigm in which mice obtain food and avoid footshocks via the same operant response. We then use this paradigm to test how the central amygdala coordinates appetitive and aversive behavioral responses. By testing somatostatin-IRES-Cre and CRF-IRES-Cre transgenic lines, we found significant differences between strains on task acquisition and performance. Using chemogenetics, we demonstrate that CeA SOM+ neurons regulate motivation for reward, while manipulation of CeA CRF+ neurons had no effect on task performance. Future studies investigating the interaction between positive and negative motivation circuits should benefit from the use of this dual valence paradigm.

## Introduction

Survival in a complex environment requires flexible responses to stimuli associated with both rewards and threats. Animal studies have revealed that several brain regions previously thought to preferentially process appetitive or aversive stimuli (e.g., amygdala, ventromedial prefrontal cortex, ventral tegmental area, cingulate cortex, periaqueductal gray) in fact respond to stimuli of either valence (Hayes et al., 2014). While there are new paradigms for simultaneous quantification of threat approach and avoidance (Heinz et al., 2017, Reis et al., 2021), few behavioral paradigms have been used that similarly assess appetitive and aversive responses (Jikomes et al., 2016, Kutlu et al., 2020). To facilitate investigation in brain regions that process oppositely valenced stimuli, we developed a paradigm to measure conditioned responses of the same modality (nose poking) to both appetitive and aversive auditory cues. This paradigm eliminates the confound of separate behavioral outputs for positive and negative reinforcement and thereby allows for direct comparison of behavioral and neuronal responses to appetitive and aversive stimuli.

We applied this novel behavioral paradigm to investigate neuronal populations in the CeA, a striatum-like structure implicated in the regulation of both defensive (Ciocchi et al., 2010; Fadok et al., 2017; Haubensak et al., 2010; Li et al., 2013; Wilensky et al., 2006) and appetitive responses (Douglass et al., 2017; Kim et al., 2017; Warlow and Berridge, 2021). The CeA modulates conditioned approach to sucrose reward (Hitchcott and Phillips,1998), and CeA lesions lead to impairment in appetitive Pavlovian conditioning (Parkinson et al., 2000) and acquisition of conditioned orienting responses (McDannald et al., 2005). Local CeA circuits generate defensive and consummatory responses through long-range projections to effector regions (Warlow and Berridge, 2021; Kong and Zweifel, 2021).

The CeA is comprised of many genetically distinct neuronal populations, and the contributions of these populations to reward and aversion are not fully understood. SOM+ and CRF+ neurons have been implicated in control of motivated behaviors. In the appetitive domain, optogenetic stimulation of either SOM+ or CRF+ neurons is positively reinforcing (Kim et al., 2017, Baumgartner et al., 2021). Additionally, pairing optogenetic stimulation of CRF+ neurons with reward delivery amplifies incentive motivation for sucrose (Baumgartner et al., 2021). Further, SOM+ neurons partially overlap with serotonin receptor 2A-expressing CeA neurons, which modulate food consumption and promote positive reinforcement by increasing perceived reward magnitude (Douglass et al., 2017). These findings indicate that CeA SOM+ and CRF+ neurons have similar roles in appetitive behaviors, although it is unclear whether these populations work synergistically or competitively during reward seeking.

SOM+ and CRF+ neurons also influence defensive and aversive behaviors. Threatening cues activate SOM+ neurons, and stimulating this population promotes freezing behavior (Li et al. 2013; Yu et al., 2016; Fadok et al., 2017). In contrast, optogenetic activation of CRF+ neurons increases anxiety-like behavior in anxiogenic contexts and promotes escape responses to threatening stimuli (Fadok et al., 2017; Paretkar and Dimitrov, 2018). These studies demonstrate that CeA SOM+ and CRF+ neurons function antagonistically to promote different threat responses.

Although SOM+ and CRF+ neurons have clear context-dependent roles in motivated behavior, natural environments are often contextually ambiguous. We therefore wished to investigate the role of CeA SOM+ and CRF+ neurons in aversive and appetitive behaviors simultaneously. We hypothesized that bidirectional chemogenetic manipulations of SOM+ and CRF+ neurons would produce similar effects in appetitive trials, specifically that performance would be improved by activation and impaired by inhibition. Additionally, we used separate appetitive tests to determine the role of these neuronal populations in the motivation to obtain reward and the drive to consume free rewards. Given the roles of the SOM+ and CRF+ populations in regulating different defensive behaviors, we hypothesized that CRF+ excitation and SOM+ inhibition would promote avoidance. Conversely, we expected that SOM+ activation and CRF+ inhibition would reduce avoidance.

## Material and methods

### Animals

Male and female C57BL/6J mice (Jackson Laboratory, Bar Harbor, ME, Stock No: 000664), heterozygous somatostatin-IRES-Cre mice (SOM-Cre; Jackson Laboratory, Bar Harbor, ME, Stock No: 028864), and heterozygous CRF-IRES-Cre mice (CRF-Cre; Jackson Laboratory, Bar Harbor, ME, Stock No: 012704) at 2-5 months old were used for the present study. Prior studies have verified high specificity of Cre expression in the extended amygdala in these lines (Partridge et al 2016; Li et al 2013). Both SOM-Cre and CRF-Cre colonies were maintained through mating with C57BL/6J mice obtained from Jackson Laboratory. Mice were individually housed on a 12 h light/dark cycle. Mice had unlimited access to drinking water but were food restricted to 85% of initial body weight. Experiments were performed during the light phase at the same time every day, at zeitgeber times (ZT) 5-10. All animal procedures were performed in accordance with the [Authors’] University animal care committee’s regulations.

### Apparatus

Experiments were conducted in standard operant conditioning chambers enclosed in sound- and light-attenuating cubicles (Med Associates, Inc., St. Albans, VT) and connected to a computer through an interface and controlled by scripts written in MED-PC V software. Each chamber was equipped with a grid floor, a house light, sound generator, two nose poke holes with tri-colored LED lights above them, and a food dispenser that delivered 20 mg food pellets (chocolate flavor, Bio-Serv, Lane Flemington, NJ) into a food receptacle located between the nose poke holes. Chambers were cleaned with 70% ethanol between subjects.

### Dual valence paradigm

#### Phase 1: Reward conditioning

The house light was illuminated during the conditioning sessions. Mice were conditioned to nose poke for food under a continuous reinforcement schedule until they reached a criterion of 50 reinforcers during a 60-min session. Tri-color LED light cues above the port indicated the active nose poke hole in each trial. These lights turned on at the beginning of each trial and turned off after the correct response (nose poke in the active nose poke port). The active port was determined randomly. New trials began immediately after the mouse entered the food receptacle to retrieve the previous reward.

#### Phase 2: Transitional phase

Each conditioning session started with 20 trials of nose poke training identical to phase 1, except that there were no light cues above the active port. Mice were required to poke in a randomized active port to get one 20 mg chocolate pellet. After this initial appetitive block, randomized appetitive and rewarded avoidance trials began. Trials began with a 30 sec auditory signal at 70 dB: either white noise or 1 kHz tone. The tone cue signaled the start of the appetitive trial; mice had 30 sec to nose poke in the active port (the side was randomized between mice and kept the same for each animal) for a pellet. If mice did not respond, a 2 sec time out period occurred, followed by the next trial. The white noise cue signaled the start of the aversive trial, during which mice had 30 sec to nose poke in a separate port to escape a footshock (1 sec, 0.2 mA). Successful avoidance resulted in pellet delivery. Failure resulted in footshock, and no reward was delivered. Successful trials were separated by a 2 sec intertrial interval. The session ended when mice earned 60 food rewards (including the initial 20 pellets at the beginning), or after 60 minutes. Mice were trained on this schedule until their footshock avoidance rate was greater than 70% or was more than 30% and stable for 2 days (<20% fluctuation).

#### Phase 3: Testing phase

The Testing phase is identical to the Transitional phase, except that successful avoidance trials do not result in pellet delivery. For chemogenetic manipulations, CNO or vehicle administration was separated by at least two sessions.

### Behavioral data collection

Behavioral data was collected automatically using Med-PC V software. The main parameters included: reinforced appetitive trials (% rewarded trials); negatively reinforced avoidance (% avoided trials), average time in seconds to correct nose pokes on appetitive and aversive trials, incorrect responses (nose poking in the opposite port) during appetitive or aversive trials. Only mice that had continuous daily training were included in the analysis of training metrics.

### Progressive ratio test

During this 60-minute test, the operant requirement for food reinforcement was 4*n, with n being the trial number. The active nose poke port was counterbalanced across animals.

### Free reward test

During this 30 min test, every head entry into the food receptacle was rewarded by a food pellet.

### Viral vectors and Surgery

For Cre-dependent chemogenetic inhibition, we used AAV-hSyn-DIO-hM4D(Gi)-mCherry (Addgene viral prep # 44362-AAV5; http://n2t.net/addgene:44362; RRID:Addgene 44362). For Cre-dependent chemogenetic excitation, we used AAV-hSyn-DIO-hM3D(Gq)-mCherry (Addgene viral prep # 44361-AAV5; http://n2t.net/addgene:44361; RRID:Addgene 44361). Control subjects were injected with AAV-hSyn-DIO-mCherry (Addgene viral prep # 50459-AAV5; http://n2t.net/addgene:50459; RRID:Addgene_50459). All vectors were used at a titer of 10^12^ particles/mL.

Viral vectors (0.3-0.5 μl) were bilaterally injected into the CeA using the following coordinates: 1.2 mm posterior and 2.85 mm lateral to the bregma, and 4.3 mm below the dura. Mice were deeply anaesthetized using 5% isoflurane (Fluriso, VetOne, Boise, ID) in oxygen-enriched air (OxyVet O2 Concentrator, Vetequip, Pleasanton, CA), followed by a subcutaneous injection of 2 mg/kg meloxicam (OstiLox, VetOne, Boise, ID), and then fixed into a stereotaxic frame (Model 1900, Kopf Instruments, Tujunga, CA) equipped with a robotic stereotaxic targeting system (Neurostar, Germany). Anesthetized mice were kept on 2-2.5% isoflurane, and a core body temperature was maintained at 36°C using a feedback-controlled DC temperature controller (ATC2000, World Precision Instruments, Sarasota, FL). Eye ointment (GenTeal, Alcon, Switzerland) was applied to the mouse’s eyes to prevent dryness. The head was shaved, and the skin was sterilized using Betadine iodine solution (Purdue Products, Stamford, CT). 2% lidocaine (0.1 ml, Lidocaine 2%, VetOne, Boise, ID) was injected subcutaneously at the site of incision and a midline incision was made with a scalpel to expose the skull. Viral vector was delivered bilaterally into CeA using pulled glass pipettes (tip diameter 10-20 μm, PC-100 puller, Narishige, Japan), connected to a pressure ejector (PDES-Pressure Application System, npi electronic equipment, Germany). Behavioral training began 7 days after surgery.

SOM- and CRF-Cre mice were assigned using blocked randomization to three experimental groups (chemogenetic inhibition, chemogenetic excitation, or control vector). Each behavioral test was repeated twice, and CNO/vehicle delivery was randomized.

For pharmacological inactivation experiments, C57Bl/6J mice were prepared for surgery as described above and bilateral stainless-steel guide cannulae (P1 Technologies) were implanted targeting the CeA. Cannulae and three stainless steel screws were affixed to the skull with Metabond, then the headcap was built up with gel superglue. Stainless steel obturators were kept in the guide cannulae until infusion.

### CNO treatment

Clozapine N-oxide (CNO; made 1 mg/ml in vehicle, given as 10 ml/kg for final dose of 10 mg/kg; Enzo Life Sciences, Farmingdale, NY) or vehicle (0.5% dimethyl sulfoxide, Sigma, St. Louis, MO, 0.9% saline, administered at 10 ml/kg volume) was injected intraperitoneally 30 min before the start of behavioral testing.

### Muscimol treatment

Muscimol (Tocris) was dissolved in 0.9% sterile saline and delivered locally into the CeA 15 minutes before behavioral testing via bilateral infusion cannulae connected to a syringe pump. A total of 400 ng/side was infused in a volume of 400 nL/side at a rate of 0.5 uL/min.

### Histology

Following testing, mice were anesthetized with tribromoethanol (240 mg/kg, i.p.) and transcardially perfused with 4% paraformaldehyde in phosphate-buffered saline (PBS). Fixed brains were cut on a Compresstome vibrating microtome (Precisionary, Greenville, NC) in 100 μm coronal slices.

Antibody staining was performed on free-floating tissue sections. After 3 × 10 min washes with 0.5% PBST, slices were blocked in 5% donkey serum in 0.5% PBST for 2 hours. Sections were incubated overnight in primary antibodies at 4°C. On the next day, sections were washed in 0.5% PBST (3 × 10 min), and then went through a 2 hr incubation with secondary antibodies at 4°C. After 3 × 10 min washes in PBS, slices were mounted using mounting medium with DAPI (Biotium, Fremont, CA). The primary antibody was rabbit anti-RFP (1:1500; 600-401-379, Rockland Immunochemicals, Pottstown, PA, RRID: AB_2209751), and the secondary antibody was goat anti-rabbit AlexaFluor555 (1:500; A-21428, Thermo Fisher Scientific, Waltham, MA, RRID: AB_2535849).

Images were obtained using an AxioScan.Z1 slide-scanning microscope (Zeiss, Germany) and a Nikon A1 Confocal microscope (Nikon, Japan). Mice were included in data analysis for Figs. 4-6 only if bilateral expression limited to the target region was observed in at least 3 consecutive brain sections (across anterior-posterior axis).

### Patch clamp electrophysiology

#### Slice preparation

Coronal brain slices containing the CeA were collected from mice at least two weeks after viral injections for *ex vivo* electrophysiological recordings. Mice were decapitated and the brains were dissected and immersed in ice-cold, oxygenated cutting solution containing (in mM): 93 N-methyl-D-glucamine, 2.5 KCl, 30 NaHCO3, 1.2 NaH2PO4, 20 HEPES, 5 Na-ascorbate; 3 Na-pyruvate, 25 glucose, 2 thiourea, 0.5 CaCl2, 10 MgSO4. The pH was adjusted to ∼7.35 with HCl. Brains were trimmed and glued to the chuck of a Leica VT-1200 vibratome (Leica Microsystems, Germany) and 300 μm-thick coronal slices were sectioned. Slices were incubated in cutting solution for 15 minutes at 34°C, then transferred to a chamber containing oxygenated artificial cerebrospinal fluid (ACSF) containing (in mM): 126 NaCl, 2.5 KCl, 1.25 NaH2PO4, 1.3 MgCl2, 2.5 CaCl2, 26 NaHCO3, and 10 glucose. Slices were maintained at 34°C for 15 min, then held at room temperature.

#### Patch clamp recording

Slices were transferred from the holding chamber to a submerged recording chamber mounted on the fixed stage of an Olympus BX51WI fluorescence microscope equipped with differential interference contrast (DIC) illumination. The slices in the recording chamber were continuously perfused at a rate of 2.5 ml/min with ACSF at 34°C and continuously aerated with 95% O2/5% CO2. Whole-cell patch clamp recordings were performed in mCherry-labeled SOM+ or CRF+ neurons in the CeL. Glass pipettes with a resistance of 3-5 MΩ were pulled from borosilicate glass (ID 1.2mm, OD 1.65mm) on a horizontal puller (Sutter P-97) and filled with an intracellular patch solution containing (in mM): 130 potassium gluconate, 10 HEPES, 10 phosphocreatine Na2, 4 Mg-ATP, 0.4 Na-GTP, 5 KCl, 0.6 EGTA; pH was adjusted to 7.25 with KOH and the solution had a final osmolarity of ∼ 290 mOsm. Series resistance was below 15 MΩ immediately after break-in and was compensated via a bridge balance circuit. To assess firing properties, 1000 ms depolarizing current injections were applied in current clamp mode. CNO (5 µM) was bath applied for a minimum of 5 minutes. Data were acquired using a Multiclamp 700B amplifier, a Digidata 1440A analog/digital interface, and pClamp 10 software (Molecular Devices, San Jose, CA). Recordings were sampled at 10 kHz and filtered at 2 kHz. Data were analyzed with Clampfit software to generate frequency response curves.

### Statistical analysis

Data were analyzed using SPSS Statistics 27 (IBM, Armonk, NY) and Prism 9 (GraphPad Software, San Diego, CA). The definition of statistical significance was *p* < 0.05. For the sake of clarity, we report the results of the interaction tests, the significant simple main effects, and the significant post-hoc tests in the main text. The results of all tests are reported in Table 1. All statistical tests were two-tailed.

#### Analysis, Figures 1 and 2

Data from C57BL/6J mice were tested for normality using the Shapiro-Wilk test and sex differences were analyzed using either an unpaired Student’s t-test or the Mann-Whitney test.

**Figure 1.**
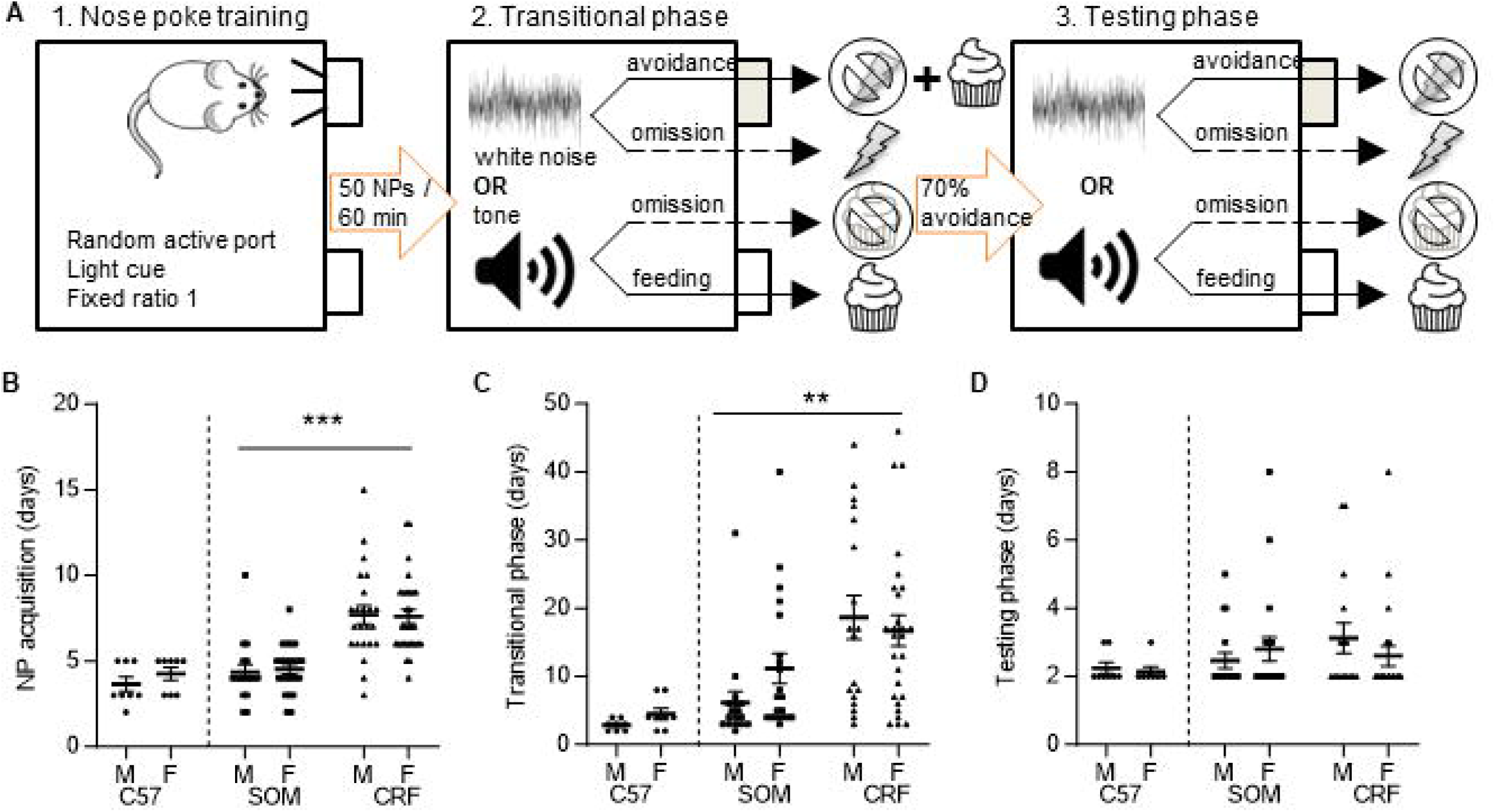
Dual valence task design and strain differences in acquisition. **A**, Overview of the three phases of the paradigm. **B**, There were no sex differences in the number of days to reach criterion for nose poke acquisition; however, CRF-Cre mice took significantly longer than SOM-Cre mice. **C**, There were no sex differences in the number of days to reach criterion in the transitional phase. CRF-Cre mice took significantly longer to acquire this phase of the task than did SOM-Cre mice. **D**, During the final phase of the task, there were no significant effects of sex or strain on the number of days to reach criteria. Data are presented as scatterplots with the mean and S.E.M. \*\**p*<0.01, \*\*\**p*<0.001

**Figure 2.**
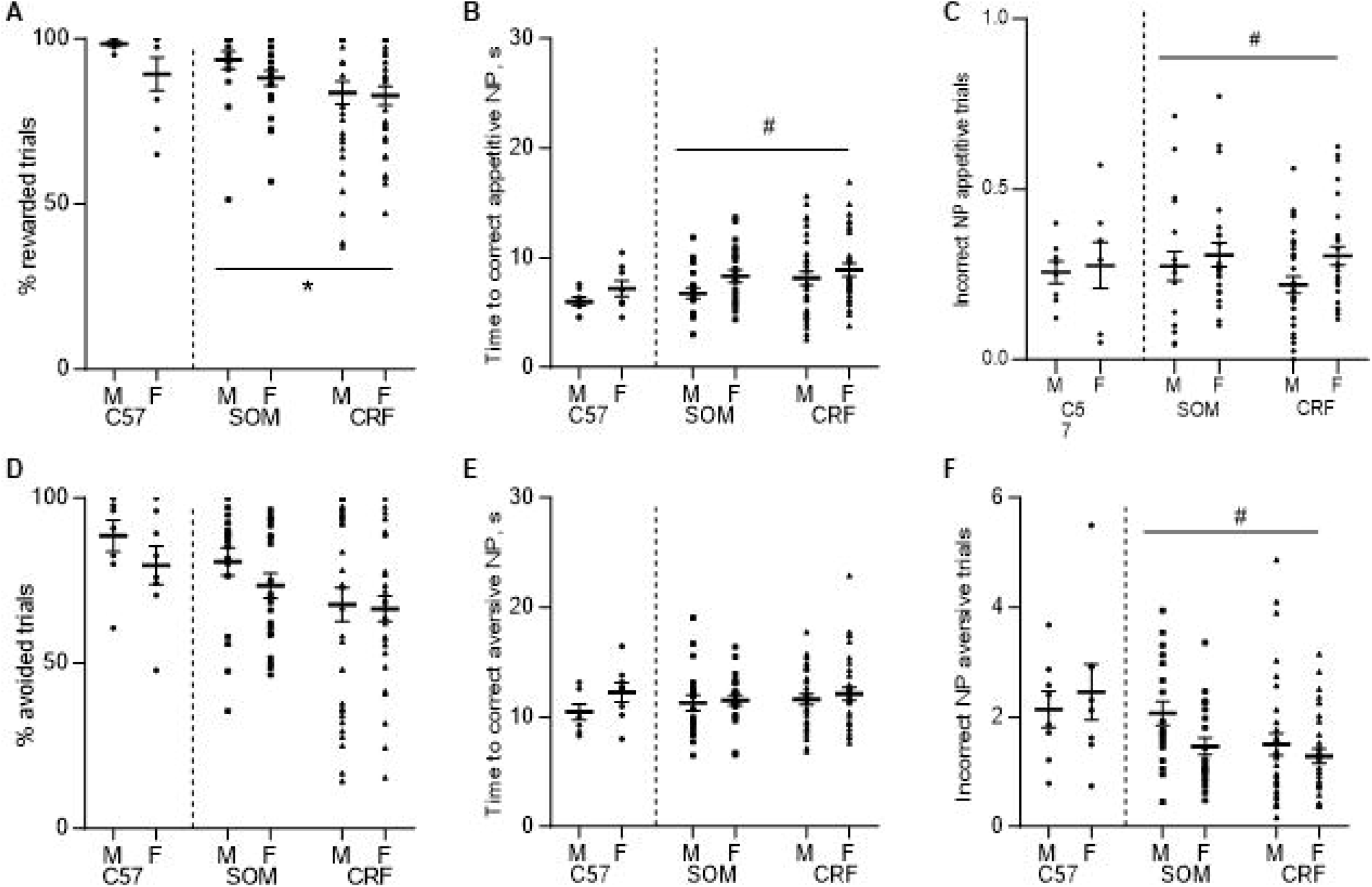
Strain and sex differences in dual valence task performance. **A**, No significant effect of sex was detected on the percentage of rewarded appetitive trials. There was a significant effect of strain, with SOM-Cre mice earning more rewards than CRF-Cre mice. **B**, Female mice took longer to make a correct response on appetitive trials. There were no strain-dependent effects. **C**, Female mice made more incorrect responses during appetitive trials. There were no strain-dependent effects. **D**, No significant effects of sex or strain were detected on successful avoidance during aversive trials. **E**, There were no significant effects of sex or strain on the latency to correct response on aversive trials. **F**, Male SOM- and CRF-Cre mice made more incorrect nose poke responses during aversive trials than did females. No significant effects of strain were detected. Data are presented as aligned dot plots with the mean and S.E.M. \**p*<0.05 (strain), #*p*<0.05 (sex) See Extended Data Figure 2-1 for the effect of intra-CeA muscimol on the dual valence task.

For strain and sex comparisons between SOM- and CRF-Cre mice, distributions of all dependent variables (DVs) exhibited skew and in some cases heterogeneity of error variance. All effects were therefore tested using generalized linear models (GLMs) analyses to model characteristics of DVs, including distribution shape, scale (continuous vs. integer-only), and whether values of zero were present. Figure 2 variables exhibiting negative skew (*% rewarded trials* and % *avoided trials*) were reverse coded to allow use of statistical models including positive skew. Reverse coding was done for significance testing purposes only and means describing significant results are reported in the DV’s original (non-reverse-coded) metric.

For Figure 1 discrete DV *Nose poke acquisition* a Poisson distribution was used in the statistical model. For DVs *Transitional phase* and *Testing phase*, skew was modeled via a negative binomial distribution as this provided better model fit than did a Poisson distribution (due to over-dispersion). For continuous DVs, gamma or Tweedie distributions were used to model skew. Figure 2 reverse-coded DV *% rewarded trials* was modeled using a Tweedie distribution, as values of zero (after reverse coding) precluded use of a gamma distribution, while % *avoided trials* was modeled using a gamma distribution. A Tweedie distribution was used in the *Incorrect NP appetitive trials* and *Time to correct aversive NP* analysis, while Gamma distributions were modeled for *Time to correct appetitive NP* and *Incorrect NP aversive trials*, because they provided better model fit than did Tweedie distributions.

#### Analysis, Figures 3-6

For Fig. 3, two-way repeated measures mixed effects analysis was applied to test the effects of current injection and CNO treatment. For Fig. 4-6, a within-subject difference score (CNO-vehicle) was calculated for each variable. Data were then tested for normality using the Shapiro-Wilk test and either an ordinary one-way ANOVA (if *p* > 0.05), or the Kruskal-Wallis test (if *p* < 0.05) was used for analysis. For Extended Data Fig. 4-1, 5-1, and 6-1, data were tested for normality using the Shapiro-Wilk test and treatment effects were analyzed using either Student’s paired t-test or the Wilcoxon test.

**Figure 3.**
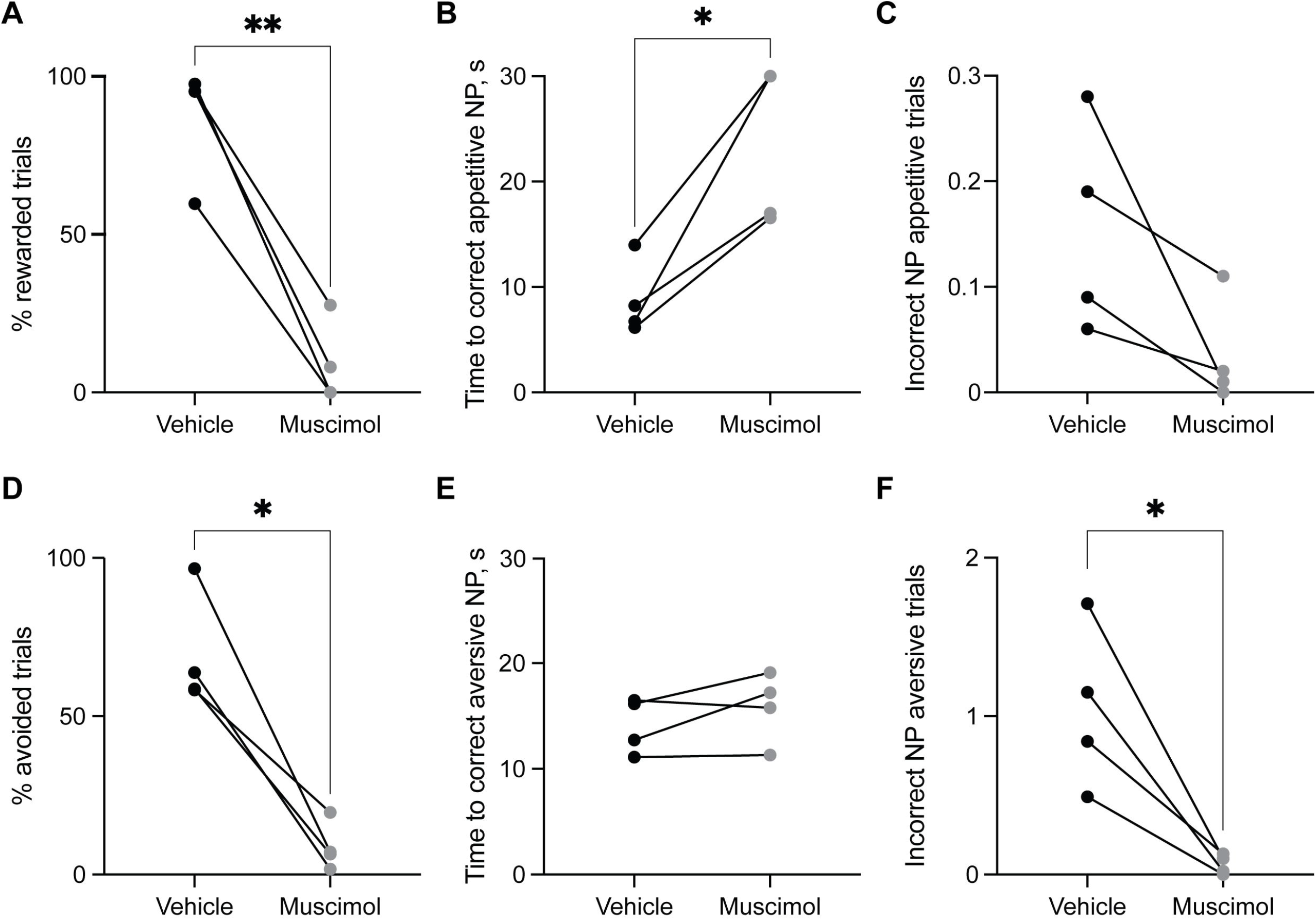
Strategy for chemogenetic manipulation of CeA SOM+ and CRF+ neurons. **A**, Three cohorts of mice per strain were injected with AAV vectors to transduce CRF or SOM neurons with either an excitatory or inhibitory DREADD. Control mice were injected with a vector expressing flurophore alone. After acquiring the dual valence task, mice were injected with CNO or vehicle 30 minutes before the task. **B**, Example images of a successful injection in a SOM-Cre mouse (top) and a CRF-Cre mouse (bottom). *Left*, bilateral expression of mCherry in the CeA. Scale bar - 2000 µm. *Right*, mCherry expression confined to the CeA. Scale bar - 1000 µm. **C,** Frequency-response relation at baseline and after treatment with 5 µM CNO in identified SOM+ neurons transfected with Gq-DREADD (left) or Gi-DREADD (right). **D,** Frequency-response relation at baseline and in CNO in identified CRF+ neurons transfected with Gq-DREADD (left) or Gi-DREADD (right). Data are presented with the mean and S.E.M. \**p*<0.05.

**Figure 4.**
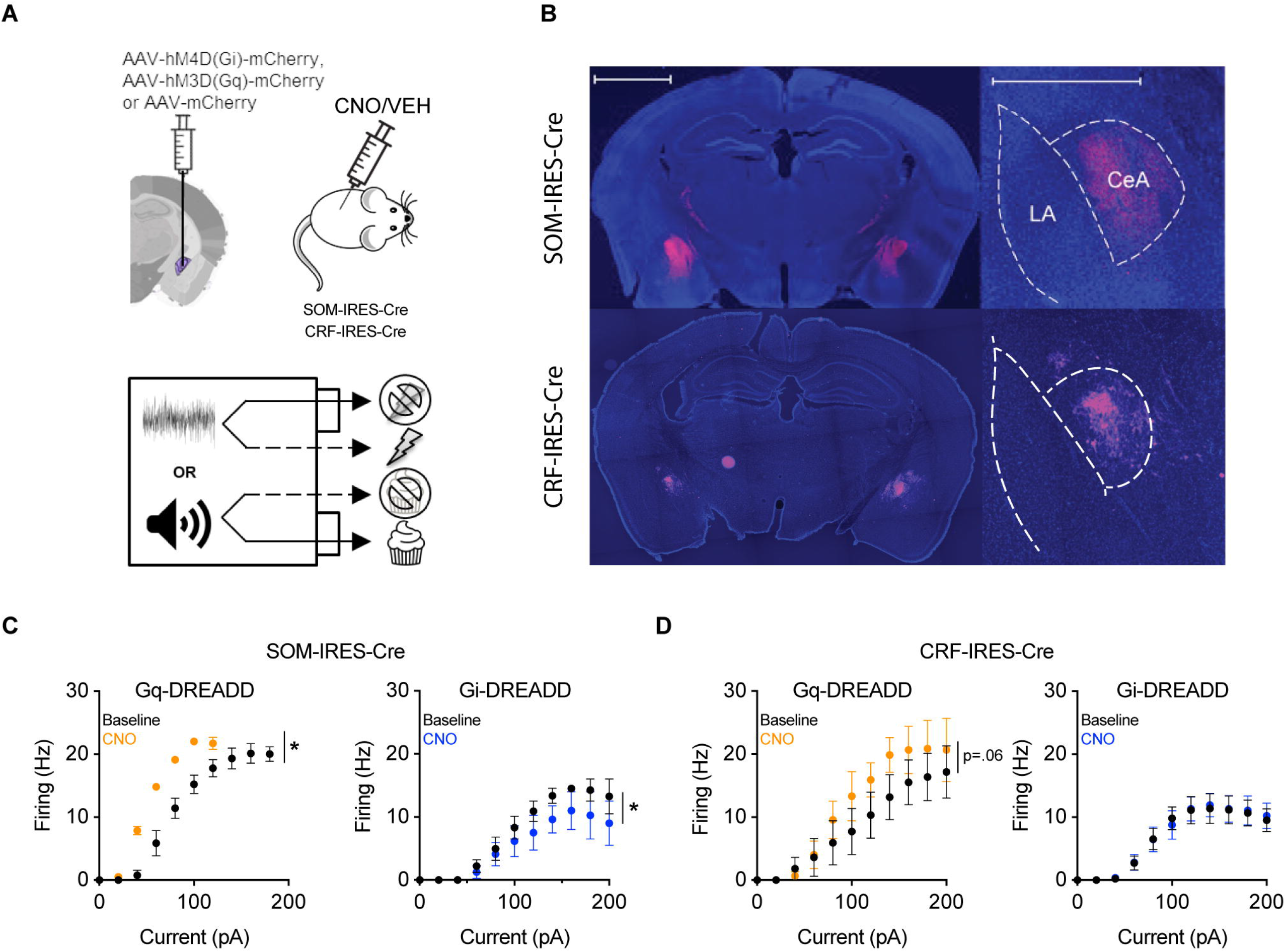
Effects of chemogenetic manipulations of CeA SOM+ neurons on task performance. A simple difference score (CNO-vehicle) was calculated for each group and performance metric. No significant between-group differences were detected for **A**, percent rewarded appetitive trials; **B**, the interval before correct response on appetitive trials; **C**, the number of incorrect nose pokes during appetitive trials; **D**, the percent of successful avoided trials; **E**, the interval before correct avoidance responses. **F**, For incorrect responses during aversive trials, a significant difference was detected between the excitatory and inhibitory DREADD groups, but neither group was significantly different than control. Box whisker plots displayed as min. to max.; boxes extend from Q1 to Q3, and horizontal lines designate the median. Triangle symbols = males, circles = females. \**p*<0.05. See Extended Data Figure 4-1 for vehicle and CNO data.

## Results

### Strain differences in acquisition of the dual valence paradigm

We developed a within-subject dual-valence operant conditioning paradigm in which mice use nose poke responses to avoid footshocks and obtain rewards in response to conditioned auditory stimuli (Fig. 1A). To test for sex differences in the acquisition of the task, equal numbers of male and female C57Bl/6J mice (N = 8 each sex) were subjected to the paradigm (Fig. 1B-D, *left*). There were no significant differences between male and female C57BL/6J mice in the number of days it took to learn the three phases of the task (Fig. 1B-D; Mann-Whitney test; NP acquisition, *U* = 23, *p* = 0.44; transitional phase, *U* = 17, *p* = 0.11; final phase, *U* = 25, *p* = 0.99). The average time needed to acquire the full task was 13±3 days.

Next, we tested for sex and strain differences in the acquisition phases of the dual valence paradigm using SOM- and CRF-Cre mice surgically prepared for chemogenetic manipulation experiments one week prior to the start of training (Fig. 1 B-D, *right*). Generalized linear models were used to analyze the effect of sex and strain on the number of days it took to reach criterion for acquisition in the three phases of the paradigm. Acquisition of the first two phases of the dual valence paradigm was significantly different between SOM- and CRF-Cre mice. Nose poke acquisition (Fig. 1B) took significantly longer in CRF-Cre (N = 23 male, 29 female) than in SOM-Cre mice (N = 17 male, 23 female; *sex X strain*, 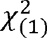 = 0.08, p= .77; main effect of *strain*, 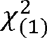 = 35.47, p < .001). The time spent learning in the transitional phase also differed significantly depending on strain (Fig. 1C). CRF-Cre mice took longer to reach criterion in the transitional phase (N = 18 male, 27 female) than did SOM-Cre mice (N = 17 male, 20 female; *sex X strain*, 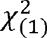 = 2.26, p = .13; main effect of strain, 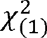 = 10.28, p = .001). There were no significant differences in the number of days it took to acquire the final phase of the task (Fig. 1D; CRF-Cre, N = 16 male, 23 female; SOM-Cre, N = 17 male, 20 female; *sex X strain*, 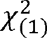 = 0.6, p= .44).

### Sex and strain differences in performance of the dual valence paradigm

To test for potential sex differences in the performance of the dual valence paradigm, we analyzed the behavior of equal numbers of male and female C57Bl/6J mice (N = 8 each sex, same mice as in Fig. 1) in the final phase of the task (Fig. 2, *left*). For appetitive trials, there were no significant differences between male and female C57BL/6J mice in the number of correct trials (Fig. 2A; Mann-Whitney, *U* = 21.5, *p* = 0.27), the latency to correct response (Fig. 2B; unpaired t-test, *t*_(14)_ = 1.4, *p* = 0.17), or in the number of responses in the opposite port (Fig. 2C; unpaired t-test, *t*_(14)_ = 0.27, *p* = 0.79). Similarly, there were no significant differences between male and female mice in the percentage of avoidance responses on aversive trials (Fig. 2D, unpaired t-test, *t*_(14)_ = 1.18, *p* = 0.26), the interval before a correct response (Fig. 2E; unpaired t-test, *t*_(14)_ = 1.56, *p* = 0.14), or in the number of responses in the opposite port (Fig. 2F; unpaired t-test, *t*_(14)_ = 0.52, *p* = 0.61).

To assess strain and sex differences in the performance of the dual valence paradigm, results of tests after vehicle injections were compared using generalized linear models for CRF-Cre (N = 29 male, 31 female) and SOM-Cre (N = 19 male, 23 female) mice prepared for chemogenetic manipulations (Fig. 2, *right*). CRF-Cre mice completed fewer successful appetitive trials than SOM-Cre mice (Fig. 2A; *sex X strain*, 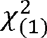 = 1.08, p = .30; main effect of strain, 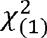 = 6.2, p = .013). A significant effect of sex was detected on the latency to correct response on appetitive trials, with female mice taking longer than males (Fig. 2B; *sex X strain*, 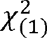 = .19, p= .66; main effect of sex, 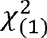 = 5.7, p = .017). Female mice also made more responses than males into the opposite port during appetitive trials (Fig. 2C.; *sex X strain*, 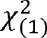 = 1.1, p= .29; main effect of sex, 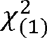 = 4.12, p = .042).

Generalized linear models were also used to analyze the effect of strain and sex on performance during avoidance trials. There were no significant differences on avoidance trial performance (Fig. 2D; *sex X strain*, 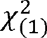 = .32, p = .574). There were also no statistically significant effects of stress or sex on the interval before a correct aversive nose poke (Fig. 2E; *sex X strain*, 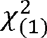 = .15, p = .702). There was, however, a significant effect of sex on the number of incorrect nose pokes on aversive trials, with males making more responses into the opposite port than females (Fig. 2F; *sex X strain*, 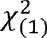 = .33, p = .568; main effect of sex, 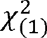 = 5.57, p = .018).

### The CeA is necessary for dual valence task performance

We next tested if the central amygdala (CeA) is necessary for performance of the dual valence task by reversibly inactivating it via local application of muscimol. C57Bl/6J mice (N = 4) with bilateral cannulae targeting the CeA were trained to criteria as in Figure 1, and muscimol (400 ng/side) or vehicle was microinjected into the CeA 15 min before testing. Vehicle and muscimol treatment occurred on nonconsecutive days, and treatment order was counterbalanced across mice. Muscimol reduced the number of rewarded trials and increased the latency to nosepoke when mice did respond for reward (Fig. 2-1 A; paired t-test, *t*_(3)_ = 8.95, *p* = 0.003; Fig. 2-1 B; paired t-test, *t*_(3)_ = 4.46, *p* = 0.021), but it did not significantly reduce the number of nose pokes in the opposite port (Fig. 2-1 C; paired t-test, *t*_(3)_ = 2.35, *p* = 0.101). On aversive trials, muscimol reduced the number of successful avoidance responses (Fig. 2-1 D; paired t-test, *t*_(3)_=5.64, *p*=0.011) without altering the latency to correct response (Fig. 2-1 E; paired t-test, *t*_(3)_=1.44, *p*=0.246) Muscimol also decreased the number of incorrect responses (Fig. 2-1 F: paired t-test, *t*_(3)_=3.99, *p*=0.028). These impairments are consistent with a role for the CeA in the performance of this dual valence task.

### Effects of CeA SOM+ chemogenetic manipulations on dual valence task performance

To determine the contribution of SOM+ and CRF+ CeA neurons to dual valence task performance, DREADD vector-injected SOM-Cre and CRF-Cre mice were injected with CNO or vehicle in two nonconsecutive sessions in a counterbalanced fashion (Fig. 3A). Following histological confirmation of targeting (Fig. 3B), data from successful cases were statistically tested. To validate the efficacy of the chemogenetic vectors, we performed patch-clamp recordings from DREADD-transfected SOM+ and CRF+ neurons. Spike frequency-response (F-I) curves were tested at baseline and in the presence of 5 uM CNO. In SOM-Cre mice, Gq-DREADD activation left-shifted the F-I relation, and Gi-DREADD activation downshifted the F-I relation (Fig. 3C; two-way repeated measures mixed model analysis, Gq CNO F_(1, 2)_ = 29.33, *p* = 0.032, *n* = 3; Gi CNO F_(1, 6)_ = 7.63, *p* = 0.033, *n* = 7). In CRF-Cre mice, Gq-DREADD activation trended towards an F-I upshift, while Gi-DREADD had no effect on the F-I relation (Fig. 3D; two-way repeated measures mixed model analysis, Gq CNO F_(1, 4)_ = 6.77, *p* = 0.060, *n* = 5; Gi CNO F_(1, 10)_ = 0.021, *p* = 0.889, *n* = 11). Given these results, we performed bidirectional chemogenetic manipulations in SOM-Cre mice, and only excitatory Gq DREADD manipulations in CRF-Cre mice.

On appetitive trials for the SOM cohorts (N = 10 mCherry, 8 Gq-DREADD, 7 Gi-DREADD), there was no significant difference between the control and DREADD groups on the effect of CNO on percentage of rewarded trials (Fig. 4A; Kruskal-Wallis test, K-W statistic = 2.5, *p* = 0.29), the interval before correct nose poke (Fig. 4B; ordinary one-way ANOVA, F_(2, 22)_ = 0.09, *p* = 0.91), or on the average number of incorrect nose pokes per trial (Fig. 4C; ordinary one-way ANOVA, F_(2, 22)_ = 3.3, *p* = 0.057). The vehicle and CNO data are presented separately for each group in Fig. 4-1 A-C.

There was no statistically significant difference detected on the effects of CNO on percent avoidance on aversive trials (Fig. 4D; ordinary one-way ANOVA, F_(2, 22)_ = 0.10, *p* = 0.90) or the time to correct nose poke (Fig. 4E; ordinary one-way ANOVA, F_(2, 22)_ = 0.58, *p* = 0.57). There was a statistically significant difference between group means on the number of incorrect nose pokes during aversive trials (Fig. 4F; ordinary one-way ANOVA, F_(2, 22)_ = 3.6, *p* = 0.043). Tukey’s multiple comparisons test found that there was a significant difference between the Gq- and Gi-DREADD groups (*p* = 0.034, 95% C.I. = [0.071, 2.0]). There was no significant difference between the control group and Gq-DREADD (*p* = 0.44) or between control and Gi-DREADD (*p* = 0.25). The vehicle and CNO data are presented separately for each group in Fig. 4-1 D-F.

### Effects of CeA CRF+ chemogenetic manipulations on dual valence task performance

We next tested for the effects of chemogenetic excitation of CeA CRF+ neurons on performance of the dual valence task. For appetitive trials, there was no significant difference between groups (N = 15 mCherry, 14 Gq-DREADD) on the effects of CNO on the percentage of rewarded appetitive trials (Fig. 5A; Mann-Whitney test, U = 73, *p* = 0.17), the time to correct response (Fig. 5B; unpaired t-test, *t*_(27)_ = 0.46, *p* = 0.65), or the average number of incorrect responses (Fig. 5C; unpaired t-test, *t*_(27)_ = 0.51, *p* = 0.61). There was also no significant between-groups effect of CNO on performance during aversive trials. There was no significant difference detected for the percentage of avoided trials (Fig. 5D; unpaired t-test, *t*_(27)_ = 0.20, *p* = 0.84), the time to correct response (Fig. 5E; unpaired t-test, *t*_(27)_ = 0.64, *p* = 0.53), or the number of incorrect responses (Fig. 5F; unpaired t-test, *t*_(27)_ = 0.60, *p* = 0.55). The vehicle and CNO data are presented separately for each group in Fig. 5-1.

**Figure 5.**
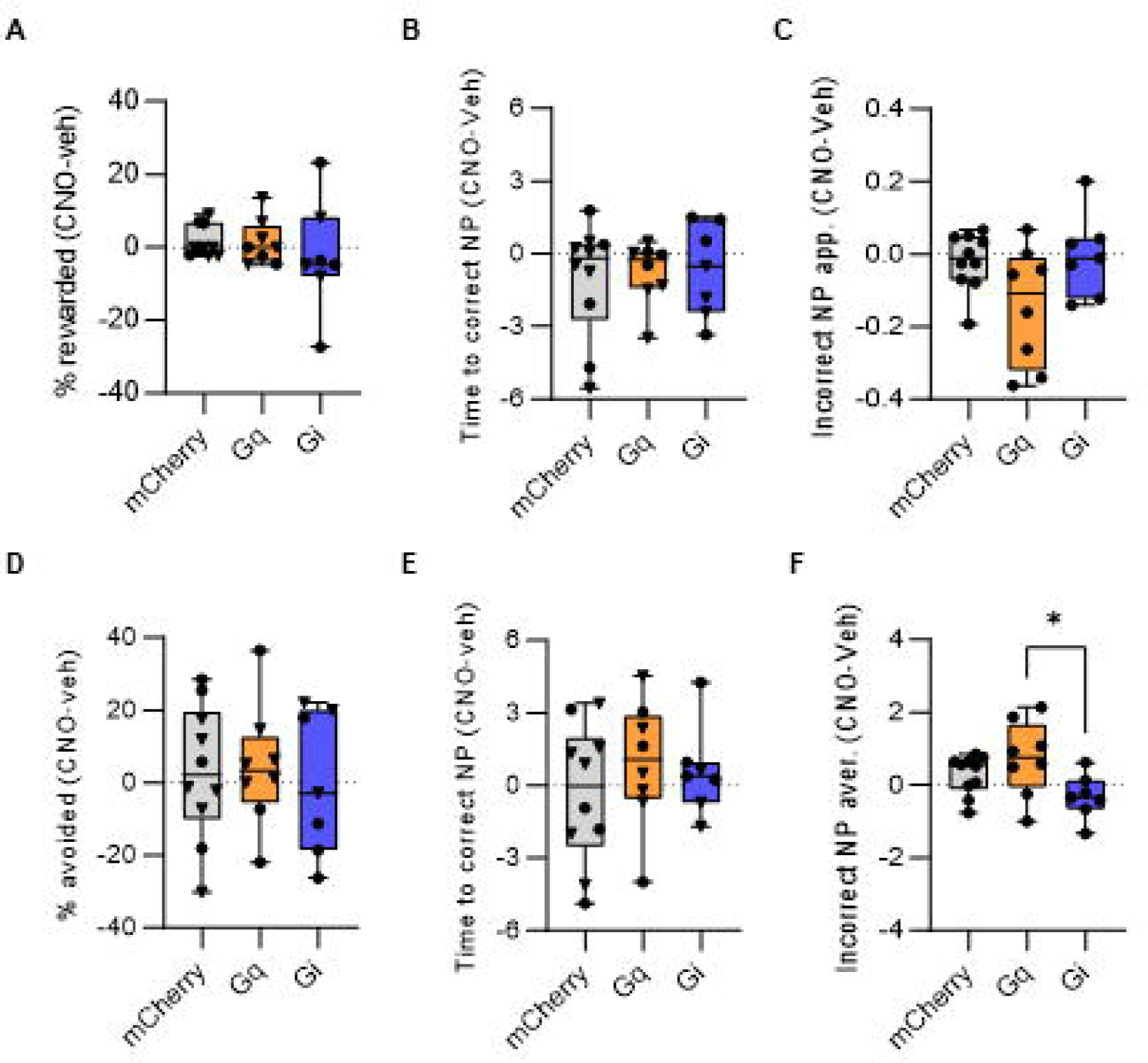
Chemogenetic manipulations of CeA CRF+ neurons has no effect on task performance. A simple difference score (CNO-vehicle) was calculated for each group and performance metric. No significant between-group differences were detected for **A**, percent rewarded appetitive trials; **B**, the interval before correct response on appetitive trials; **C**, the number of incorrect nose pokes during appetitive trials; **D**, the percent of successful avoided trials; **E**, the interval before correct avoidance responses; **F**, Incorrect responses during aversive trials. Box whisker plots displayed as min. to max.; boxes extend from Q1 to Q3, and horizontal lines designate the median. Triangle symbols = males, circles = females. See Extended Data Figure 5-1 for vehicle and CNO data.

**Figure 6.**
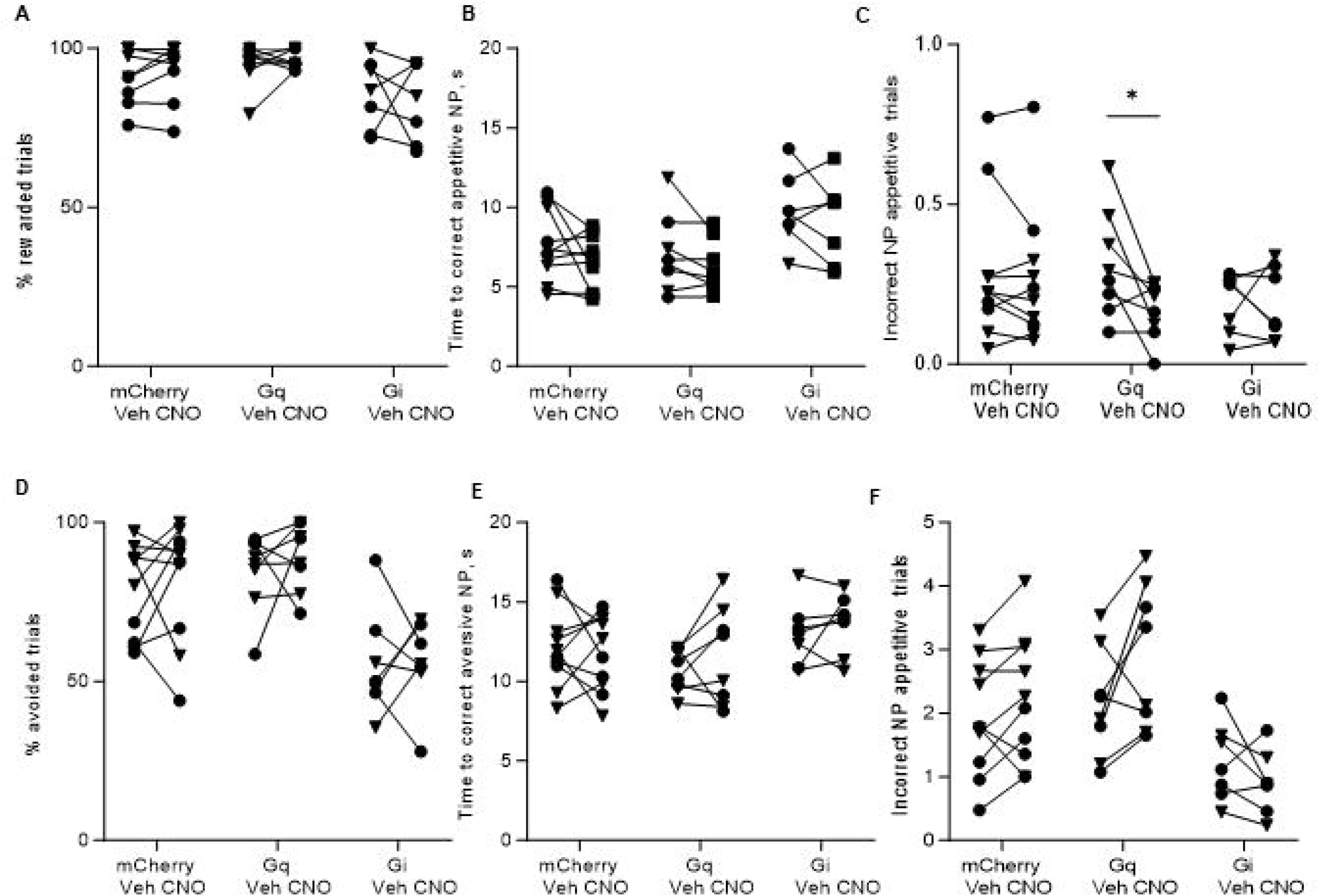
Effects of chemogenetic manipulations on appetitive motivation and free reward consumption. **A**, Chemogenetic inhibition of CeA SOM+ neurons significantly increased appetitive motivation. **B**, There were no significant differences between groups in free reward consumption with chemogenetic manipulations of SOM+ neurons. **C**, Chemogenetic manipulations of CeA CRF+ neuronal function had no effect on progressive ratio performance. **D**, There were no significant differences in free reward consumption between the CRF-Cre groups. Box whisker plots displayed as min. to max.; boxes extend from Q1 to Q3, and horizontal lines designate the median. Triangle symbols = males, circles = females. \*\**p*<0.01. See Extended Data Figure 6-1 for vehicle and CNO data.

### Chemogenetic manipulations of CeA SOM+ and CRF+ neurons during appetitive tests

In addition to understanding the effects of chemogenetic manipulations of CeA SOM+ and CRF+ neurons on performance in the dual valence task, we also sought to test the effects of these manipulations on appetitive motivation and appetite. Therefore, we compared the effects of CNO injection between groups during a progressive ratio session and a free reward consumption session (Fig. 6 and Fig. 6-1).

In SOM-Cre mice (N = 10 mCherry, 8 Gq-DREADD, 7 Gi-DREADD), a significant difference was detected between groups during the progressive ratio test (Fig. 6A; ordinary one-way ANOVA, F_(2, 22)_ = 7.2, *p* = 0.0038). Tukey’s multiple comparisons test found that there was a significant difference between the mCherry control and the Gi-DREADD groups (*p* = 0.0028, 95% C.I. = [−13, −2.7]), with CNO increasing the number of reinforcements in the Gi-DREADD group. There was no significant difference between the control and Gq-DREADD group (*p* = 0.16) or between the Gq- and Gi-DREADD groups (*p* = 0.17). There was no significant difference detected between groups on the effect of CNO on free reward consumption (Fig. 6B; ordinary one-way ANOVA, F_(2, 22)_ = 1.5, *p* = 0.25).

No significant difference was detected between the CRF-Cre groups (N = 14 mCherry, 13 Gq-DREADD) during the progressive ratio test (Fig. 6C; unpaired t-test, *t*_(25)_ = 0.94, *p* = 0.36). There was also no significant difference between groups in the effect of CNO injection during the free reward session (Fig. 6D; unpaired t-test, *t*_(25)_ = 1.7, *p* = 0.09).

## Discussion

We present a novel operant conditioning paradigm that allows measurement of approach and avoidance behaviors within a single session using an identical operant response, with similarly robust responding in appetitive and aversive trials. This paradigm simultaneously assesses numerous behavioral measures including operant performance, response latency, and incorrect perseverative responses, across valences in a single context. Importantly, by eliminating the confound of separate operant response modalities, this paradigm allows for direct comparison of the effects of genetically targeted manipulations on positive and negative reinforcement.

Cre-recombinase driver mouse lines are widely used for genetically targeted optogenetic and chemogenetic manipulations of neuronal activity. Our study revealed that heterozygous CRF-Cre mice showed a substantial delay in acquisition of operant reward and avoidance relative to C57Bl/6J and heterozygous SOM-Cre mice, another C57Bl/6J congenic line. A limitation of the dual valence paradigm is that mice requiring prolonged training in the reward conditioning or transitional phases risk appetitive overtraining, which is known to affect measures of cognitive flexibility (Caglayan et al., 2021; Garner et al., 2006). The speed of initial appetitive learning may therefore influence learning of the transitional phase, which requires cognitive flexibility. Likewise, mice requiring prolonged training in transitional and/or testing phases experience greater cumulative footshock exposure, which may induce confounding stress effects on motivated behavior (Conrad, 2010; Dieterich et al., 2021), although chronic irregular mild footshock has been shown to induce behavioral changes distinct from other chronic stress models, such as hyperactivity or changes in consumption of palatable food (Cao et al., 2007). As strain differences in acquisition of appetitive reinforcement and avoidance have been observed previously (Padeh et al 1974; Ingram & Sprott 2013), we urge caution in interpreting results from strains that do not readily acquire the dual valence task.

Recent studies have illuminated sex differences in mouse behavioral strategies in response to aversive stimuli (Keiser et al 2017; Borkar et al 2020). Studies examining sex-dependent effects on acquisition and performance of appetitive and aversively motivated operant responding in adult mice have yielded conflicting results (Padeh et al 1974; Mishima et al 1986; Kutlu et al 2020). We therefore compared acquisition and performance in the dual valence paradigm in male and female mice. We observed that female mice took longer to make a correct appetitive nose poke, made more incorrect responses during appetitive trials, and made fewer incorrect responses during avoidance trials. This effect is unlikely to result from sex differences in cognitive flexibility (switching from reward-seeking to avoidance), as prior work has found comparable cognitive performance in both sexes (Bissonette et al., 2012). Rather, this may reflect sex differences in cue discrimination (Rodríguez et al., 2011), with a bias towards the aversive cue.

Previous studies have linked CeA SOM+ and CRF+ neurons to both appetitive and aversive motivation and behaviors (Ciocchi et al., 2010; Douglass et al., 2017; Fadok et al., 2017; Haubensak et al., 2010; Kim et al., 2017; Li et al., 2013; Warlow and Berridge, 2021; Wilensky et al., 2006). Therefore, we hypothesized that chemogenetic manipulations of these neuronal populations would alter performance in the dual valence task. We were unable to determine the effect of chemogenetic inhibition of CRF+ neurons because we could not validate inhibition *in vitro*. Contrary to our hypothesis, excitation of CRF+ neurons did not significantly affect task performance when compared to control. One explanation for this negative result could be that CRF-Cre mice require significantly longer to acquire the task, potentially leading to overtraining thereby minimizing the importance of this cell type for task performance. It is possible that CRF neurons play a role in the acquisition of the task, and this could be tested in future studies.

The results of the SOM manipulations are more puzzling, given that the SOM-Cre line readily acquires the task at a similar rate to C57Bl6/J mice. The CeA SOM+ population includes food-responsive cells (Ponserre et al., 2022), and excitation of CeA SOM+ neurons projecting to the lateral substantia nigra has been shown to induce intracranial self-stimulation and real-time place preference. At the same time, inhibition of this population did not disrupt performance (Steinberg et al., 2020). Silencing of CeA SOM+ neurons has been shown to lead to impaired fear learning, while activation of these neurons sufficiently induced unconditioned and conditioned defensive behaviors (Li et al. 2013; Fadok et al. 2017; Kong & Zweifel, 2021), which we did not observe in this paradigm.

The results of the appetitive tests demonstrate that inhibition of CeA SOM+ neurons induces a significant increase in motivation to nose poke for a food reward. These results conflict with previous studies supporting a role for SOM+ CeA neurons in positive reinforcement (Douglass et al., 2017; Kim et al., 2017). It is possible that when mice are in more complex environments, SOM+ neurons are biased more toward generating negative valence behavior, or that the role of SOM+ neurons in generating consummatory behavior is altered by experience and extended learning. Alternatively, chemogenetic inhibition of SOM+ CeA neurons may alter the state of parallel CeA networks mediating feeding (Barbier et al 2020).

In conclusion, although chemogenetic manipulations of CeA CRF+ and SOM+ neurons did not elicit the hypothesized performance differences, muscimol-mediated inactivation of the CeA did dampen multiple performance metrics indicating that the dual valence paradigm we present can be used to explore the neuronal mechanisms influencing distinct types of reinforcement. For example, given that heterogeneity within the CRF+ or SOM+ CeA populations, based on localization within the CeA, or by projection targets, is important for controlling different valenced behaviors, future studies incorporating intersectional viral vector strategies are warranted.

## Acknowledgments

This work was supported by the Louisiana Board of Regents through the Board of Regents Support Fund (LEQSF(2018-21)-RD-A-17) and the National Institute of Mental Health of the National Institutes of Health under award number R01MH122561 to JPF. The content is solely the responsibility of the authors and does not necessarily represent the official views of the National Institutes of Health. MD is currently affiliated with Institute of Developmental Neurophysiology, Center for Molecular Neurobiology, University Medical Center Hamburg-Eppendorf, Hamburg, Germany.

## Animal use statement

All animal procedures were performed following Tulane University animal care committee’s regulations.

---

**Figure 2-1.**
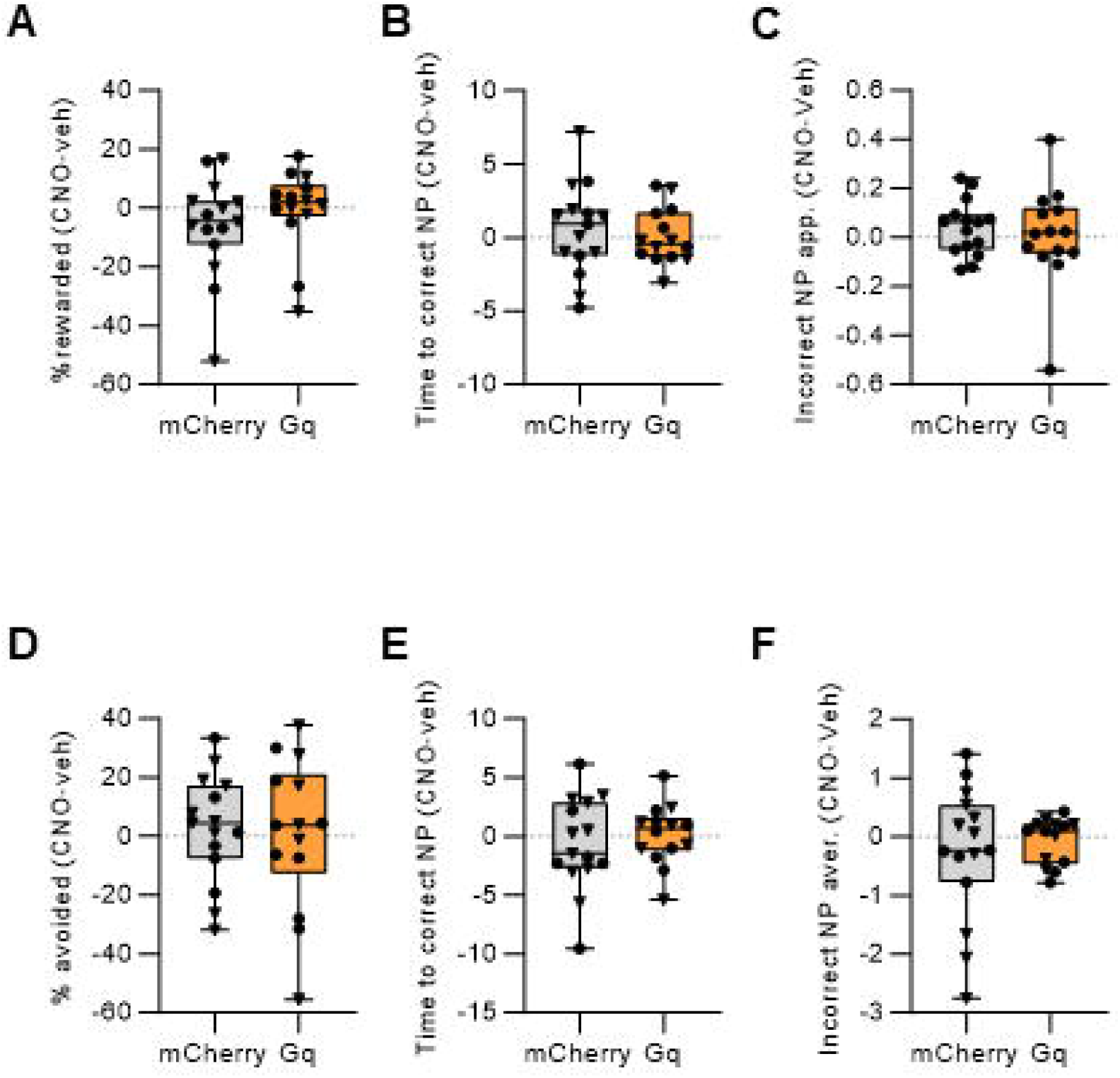
Dual valence task performance requires the CeA. Mice were implanted with bilateral cannulae targeting the CeA and muscimol (400 ng/side) or vehicle was infused prior to testing. **A**, Muscimol treatment significantly impaired appetitive operant performance. **B**, Muscimol treatment significantly increased the latency to correct response on appetitive trials. Two mice did not respond on any appetitive trials, so latency was capped at the trial duration (30 s). **C**, Muscimol caused a non-significant decrease in the average number of incorrect responses during appetitive trials. **D**, Muscimol treatment significantly impaired operant performance on avoidance trials. **E**, The latency to correct response on aversive trials was not affected by muscimol. **F**, Muscimol caused a non-signficicant decrease in the average number of incorrect responses during aversive trials.

**Figure 4-1.**
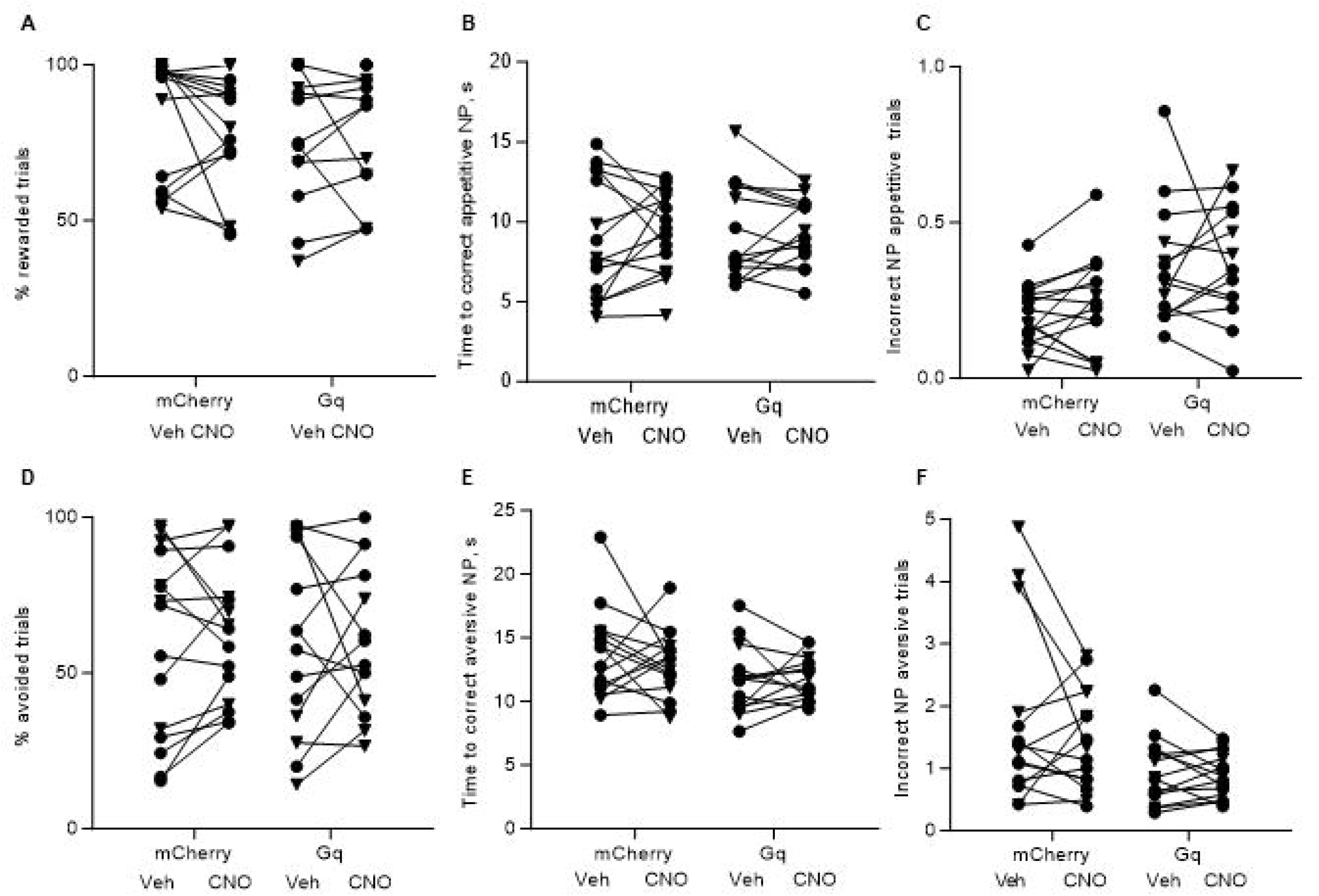
Vehicle and CNO data for the SOM-Cre chemogenetic groups. **A**, There were no significant differences between vehicle and CNO treatments on the percentage of rewarded trials . **B**, There were no significant treatment effects on the latency to correct response on appetitive trials. **C**, In the Gq group, CNO treatment caused a significant reduction in the number of incorrect responses during appetitive trials (paired t-test, *t*_(7)_ = 2.5, *p* = 0.04). **D**, There were no significant differences between vehicle and CNO treatments on the percentage of correct avoidance trials. **E**, There were no significant effects of CNO on the latency to correct avoidance response. **F**, There were no significant effects of CNO on the number of incorrect responses during aversive trials. \**p*<0.05. Triangle symbols = males, circles = females.

**Figure 5-1.**
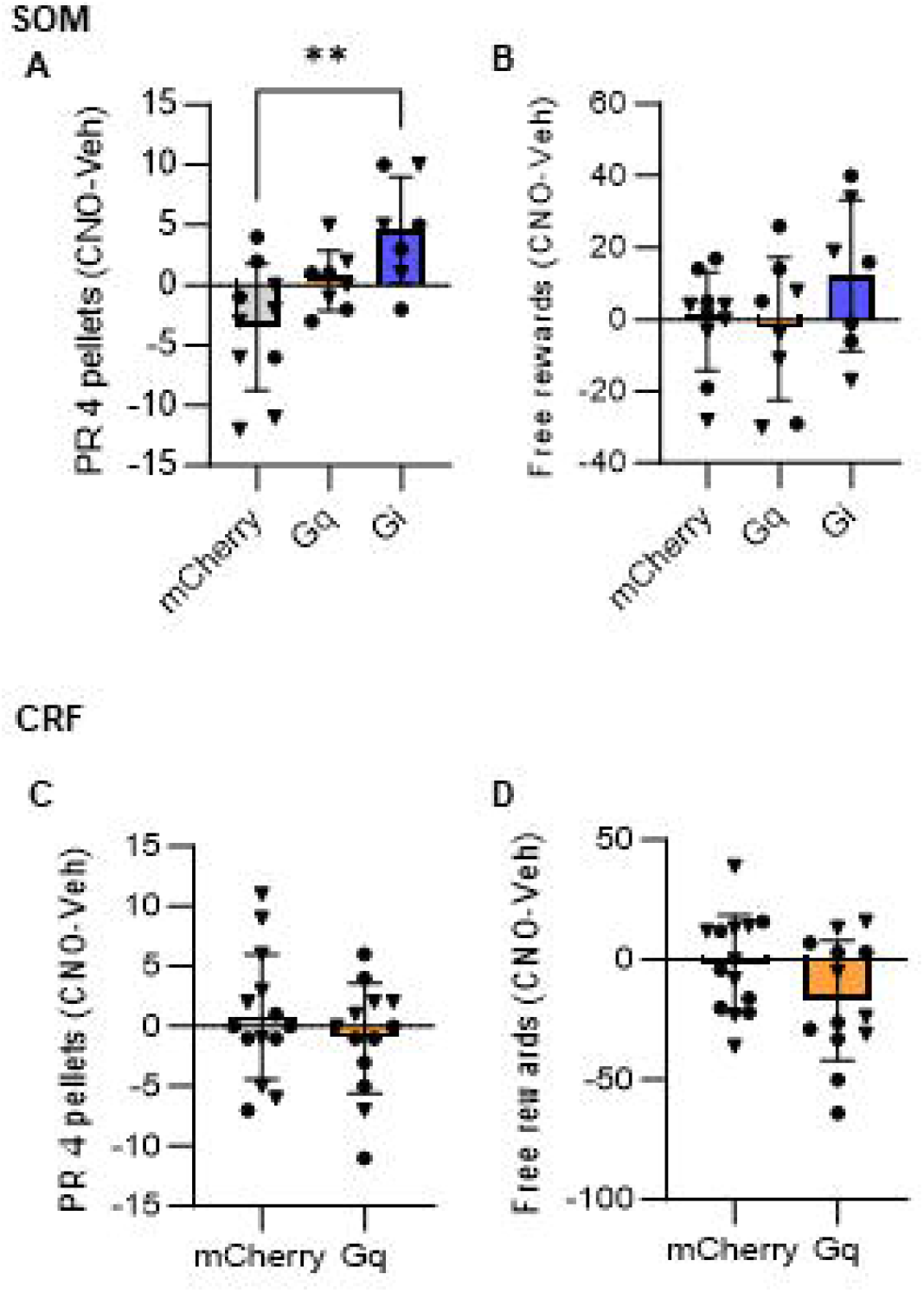
Vehicle and CNO data for the CRF-Cre chemogenetic groups. There were no significant differences between vehicle and CNO treatments on **A**, the percent of rewarded appetitive trials.**B**, the latency to correct response on appetitive trials. **C**, the number of incorrect responses during appetitive trials. **D**, percent avoidance. **E**, the latency to correct avoidance response. **F**, the number of incorrect responses during aversive trials.

**Figure 6-1.**
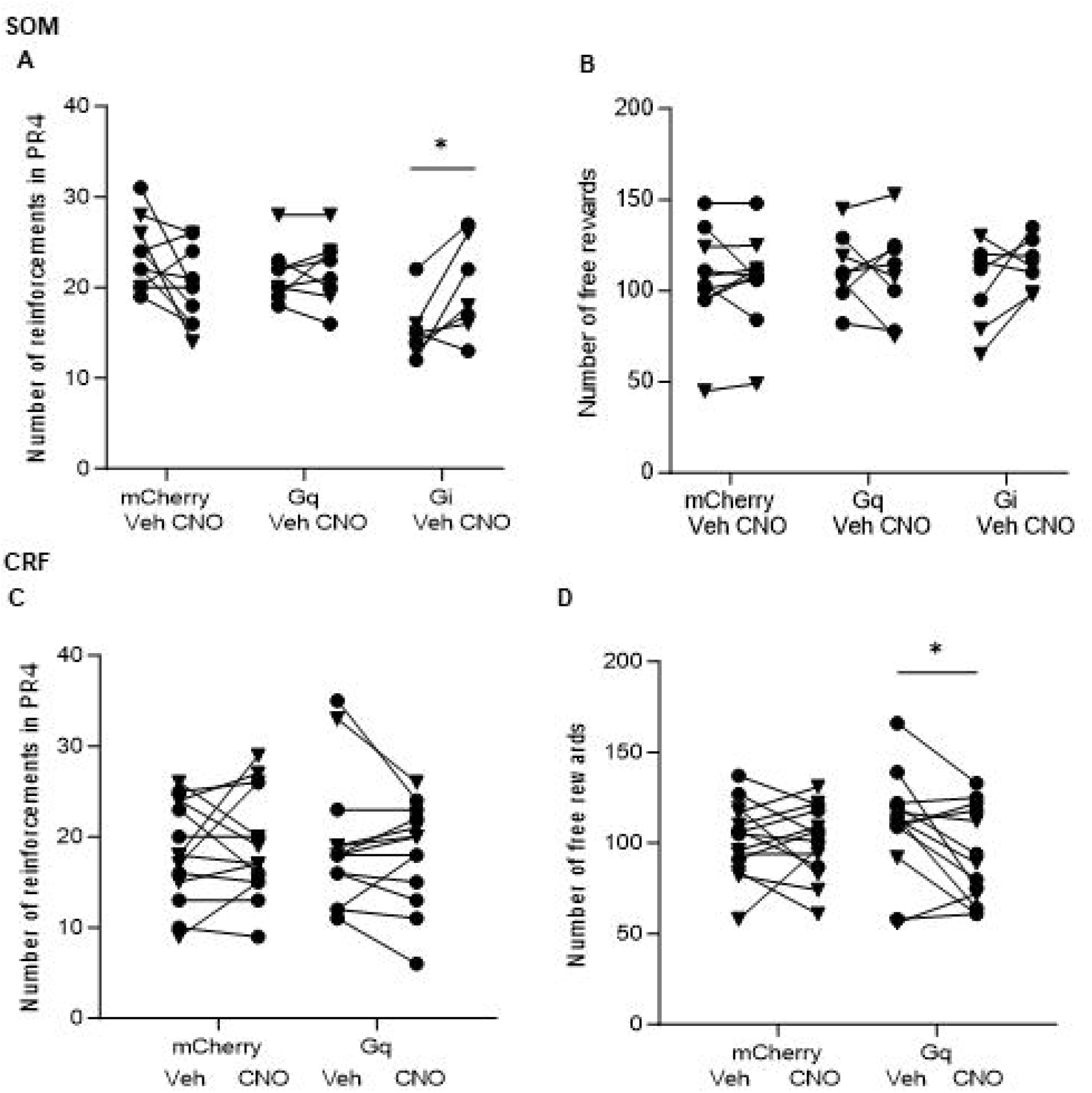
Vehicle and CNO data for the appetitive motivation and free reward consumption tests. **A**, CNO induced a significant elevation in the number of reinforcements during the progressive ratio test in the inhibitory DREADD group (paired t-test, *t*_(6)_ = 2.7, *p* = 0.03). **B**, There was no significant effect of CNO on free reward consumption in the SOM+ groups. **C**, There was no significant effect of CNO on appetitive motivation in the CRF+ groups. **D**, CNO reduced free reward consumption in the excitatory DREADD CRF group (paired t-test, *t*_(12)_ = 2.4, *p* = 0.03). \**p*<0.05. Triangle symbols = males, circles = females.

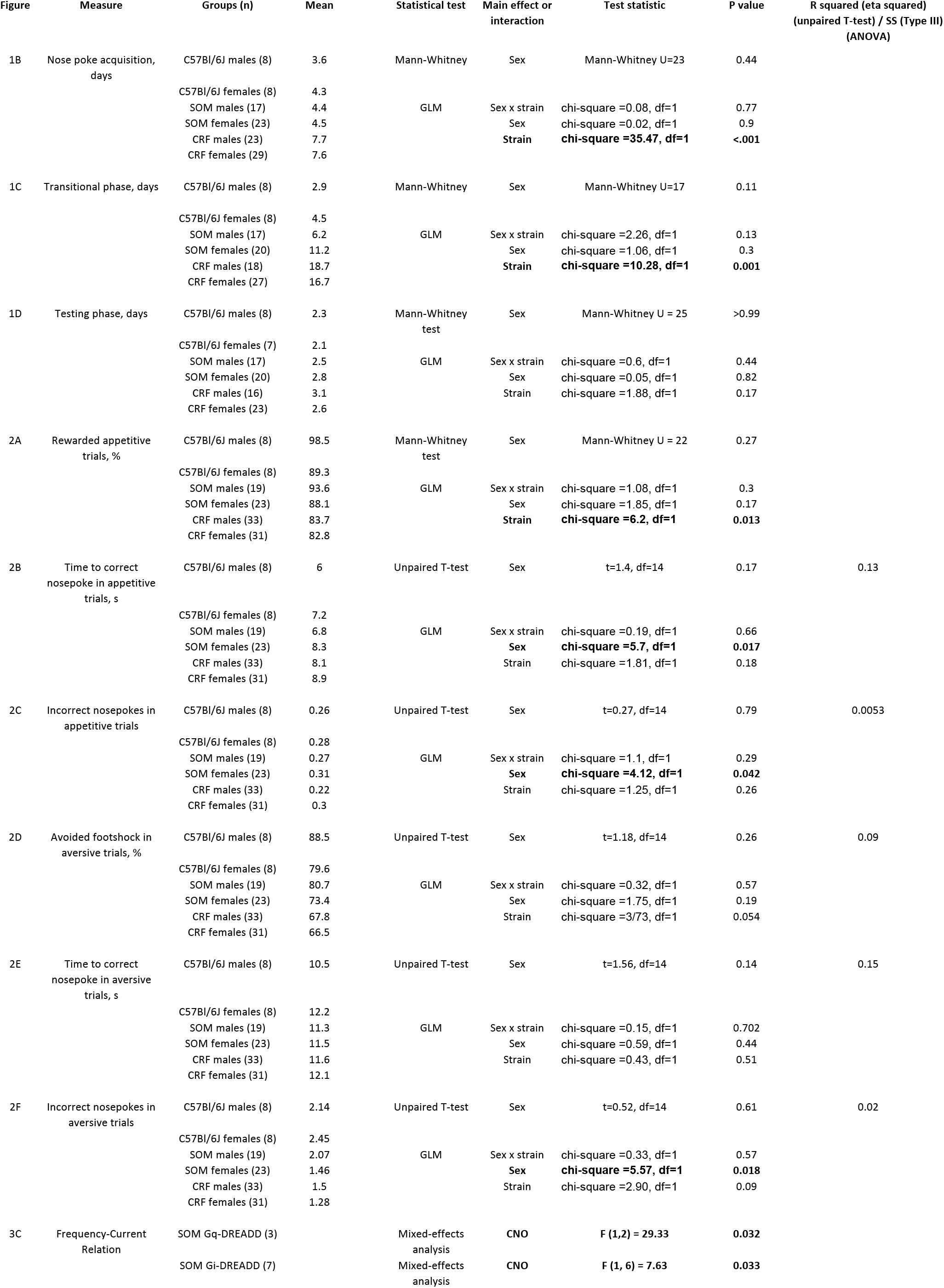
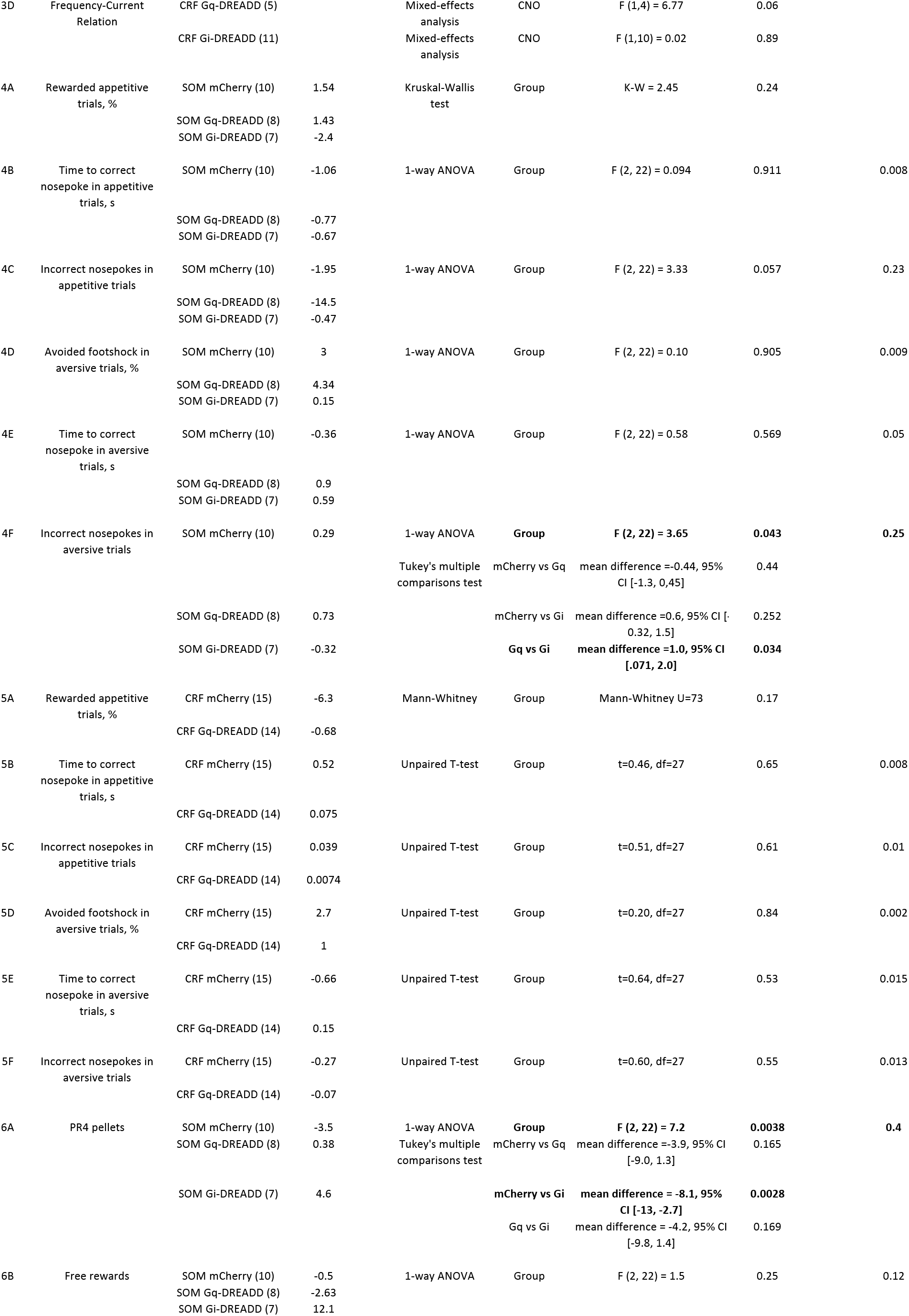
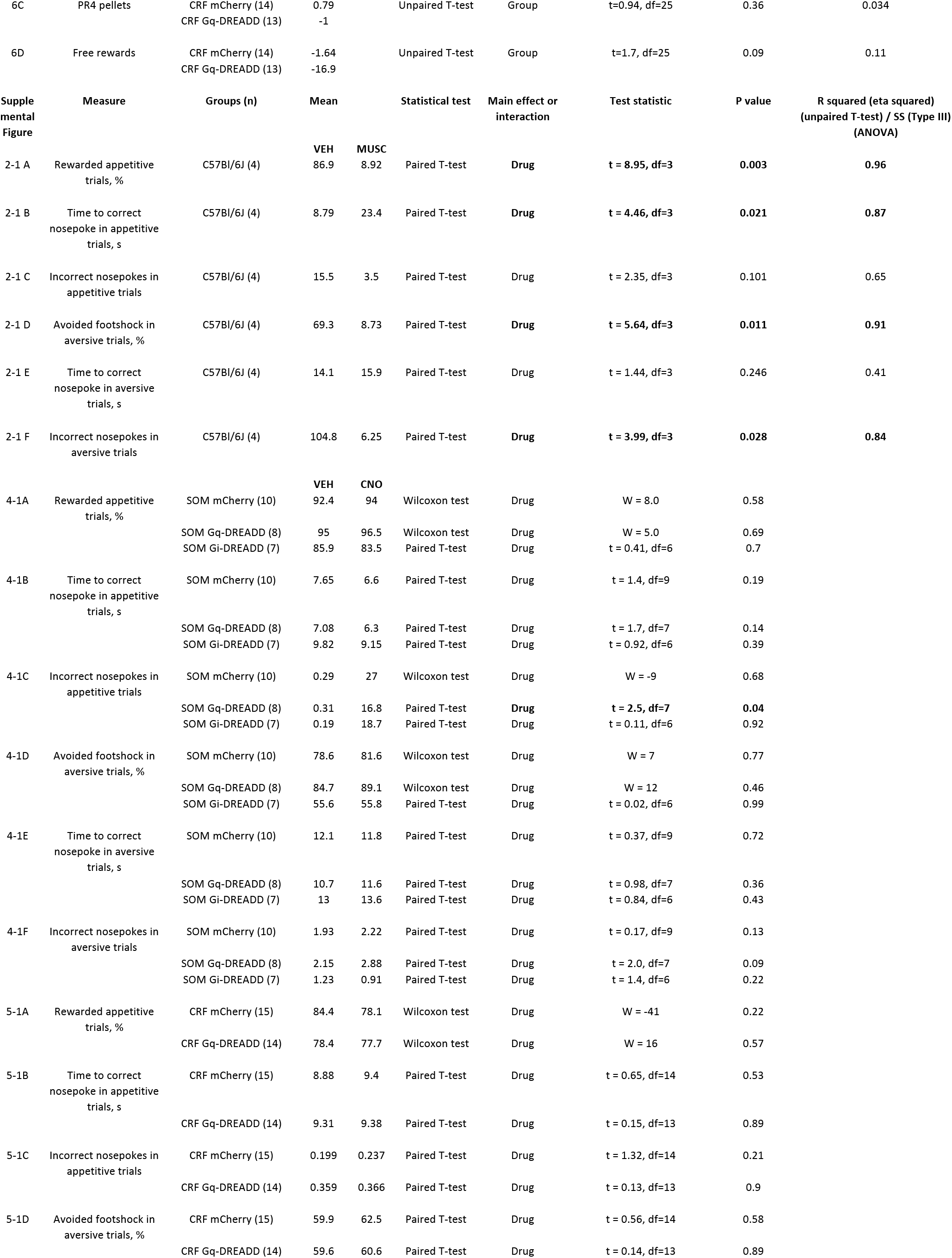
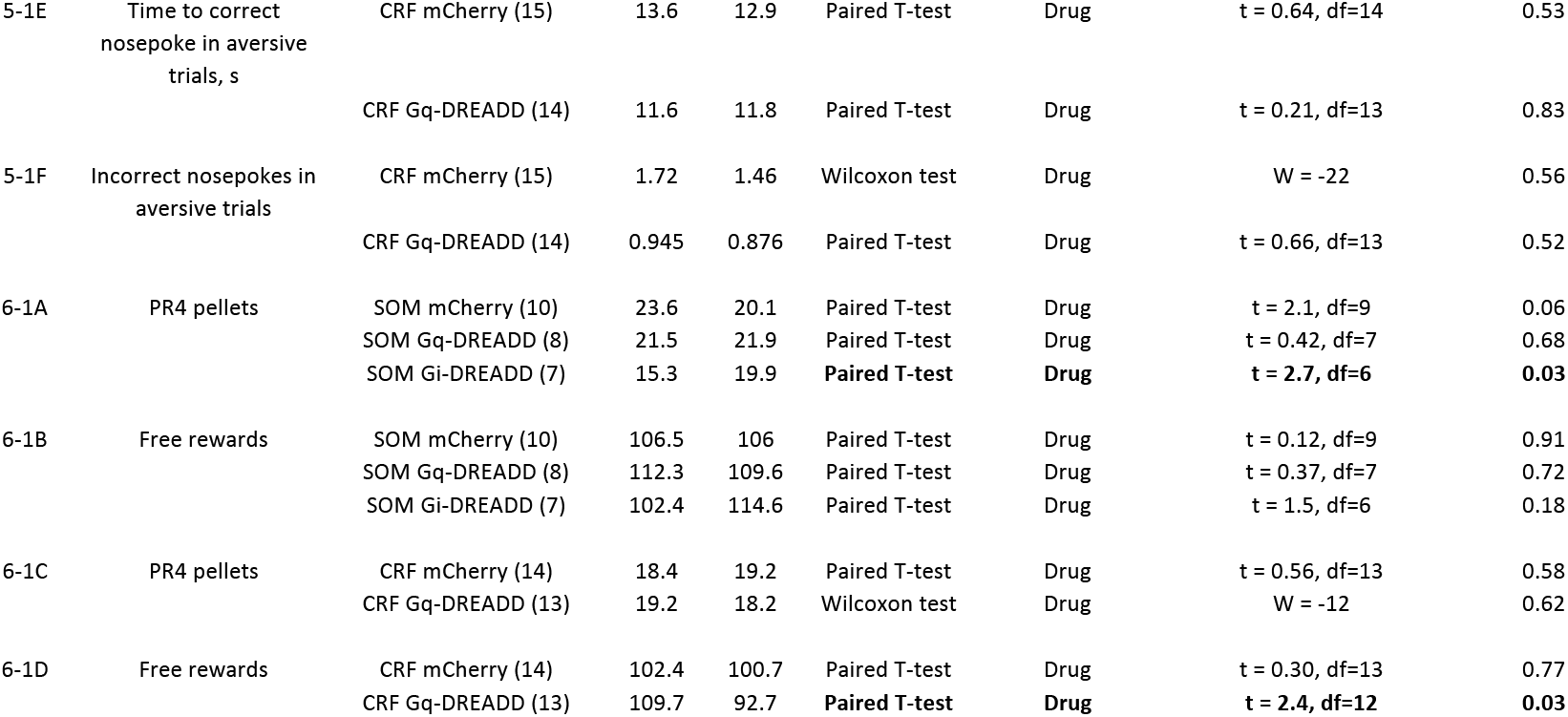

